# Longitudinal dynamics of ethanol-resistant microbes and sporulation in the human gut

**DOI:** 10.1101/2025.07.08.663423

**Authors:** Ursa Miklavcic, Aleksander Mahnic, William R. Shoemaker, Jacopo Grilli, Sandra Janezic, Maja Rupnik

**Affiliations:** Department for Microbiological Research, National Laboratory for Health, Environment and Food (NLZOH), Maribor, Slovenia; Biotechnical Faculty, University of Ljubljana, Ljubljana, Slovenia; Faculty of Medicine, University of Maribor, Maribor, Slovenia; Quantitative Life Sciences, The Abdus Salam International Centre for Theoretical Physics (ICTP), Trieste, Italy

**Keywords:** sporobiota, microbiota, longitudinal study, ethanol-resistant bacteria, community-level, sporulation frequency

## Abstract

**Background:** A considerable proportion of human gut bacteria use some form of dormancy to survive unfavorable conditions such as oxygen exposure, antibiotic treatment, and inflammation. Dormancy strategies enable persistence, allow recolonization following disruptions, and facilitate transmission between individuals. In this study, we explored the composition, characteristics, and longitudinal dynamics of the dormant community in the gut as determined by ethanol resistance. We conducted a six-month longitudinal study on nine healthy male adults, who were sampled every two weeks. Gut microbiota composition was determined with 16S rRNA amplicon sequencing in untreated (bulk microbiota) and ethanol treated samples. A rigorous algorithm was implemented to determine the ethanol-resistant and non-resistant community within the gut microbiota.

**Results:** We detected ethanol-resistant gut bacteria in eight distinct phyla, with the highest number of OTUs observed in the phylum *Bacillota*, and substantial representation in *Actinomycetota* and *Pseudomonadota*. In general, ethanol-resistant taxa exhibited higher relative abundance and longer within-individual persistence compared to non-resistant taxa. Ethanol resistance indicated a higher rate of sharing between individuals, independent of phylum. Ethanol-resistant *Bacillota* also exhibited longer persistence within the gut over the six-months period in comparison to ethanol non-resistant *Bacillota* and other ethanol-resistant or ethanol non-resistant phyla. Finally, the analysis of sporulation frequency of highly prevalent ethanol-resistant *Bacillota* members revealed that sporulation occurred in each individual synchronously among different community members but varied significantly between individuals. Major shifts in sporulation frequency were associated with well-known disturbances such as antibiotic therapy and travel.

**Conclusion:** Our findings highlight that sporulation of *Bacillota* members and alternative dormancy strategies employed by non-*Bacillota* members contribute to bacterial persistence in the human gut and facilitate transmission between unrelated individuals. Moreover, we show that sporulation frequency (specific to *Bacillota*) is a dynamic, community-level process shaped by host- and time-specific environmental factors rather than intrinsic, species-specific traits.

## Background

The composition of the human gut microbiota is a result of constant immigration and competition between bacteria, viruses, archaea, and eukaryotes. Colonization begins at birth, when infants acquire bacteria from their caretakers and environment [1]. Successful colonization of commensal bacteria depends on their capacity to spread between humans and their ability to establish and grow within them. The likelihood of transmission increases when bacteria utilize different life-strategies, including sporulation, aerotolerance, and the viable but non-culturable state to survive intermediate reservoirs and successfully colonize a new host [2]. However, high within-individual strain abundance is typically not driven by transmission, but rather by its ability to adapt and grow in the human gut [3].

Sporulation is one of the most researched strategies that increases survival time outside the human gut. At least 50% of gut bacteria possess the genetic capability for sporulation. All currently known spore-forming bacteria belong to the phylum *Bacillota*, which is a highly abundant and prevalent phylum in the human gut [4–7]. The ability of spore-forming bacteria to endure unfavorable conditions, such as oxygen, antibiotic treatment, and inflammation, allows them to persist and potentially recolonize the gut following disruptions, but also facilitates transmission between individuals [8,9].

Most studies have focused on pathogenic spore formers related to food safety and intestinal infections [10]. Far less is known about the ecological dynamics of commensal spore-forming bacteria, which are highly abundant in healthy individuals [5,9]. Up-to-date, only two culture-independent studies employed either lysozyme treatment or ethanol shock to remove vegetative cells to investigate the transmission of spore-forming bacteria between humans [11,12]. In a cohort of healthy human adults, OTUs found in the lysozyme-resistant community were more frequently shared between individuals than other taxa [11]. Furthermore, a comparison of infant-mother pairs showed that the majority of OTUs determined to be spore-forming were acquired from the environment rather than mother-to-child transmission [12]. Spore-forming bacteria are present at lower abundance in the infant gut compared to adults, but they still play an important role as early butyrate producers [13–15]. An analysis of multiple *Bacillota* genomes from the human gut revealed that the loss of sporulation ability increased the relative abundance of a given bacterial species within an individual, but decreased its overall prevalence in the broader population [16]. The persistence of spore-forming bacteria in the human gut was experimentally assessed by monitoring a single individual for a period of 24 days, revealing a stronger correlation among OTUs in the lysozyme-resistant community compared to other OTUs [11]. These findings align with the ecological concept of a microbial seed bank, where dormant life stages, such as spores, form a reservoir of potential colonizers capable of repopulating the community following disturbance [17,18].

In this study, we analyzed the longitudinal dynamics of the ethanol-resistant bacterial community in the human gut, with a specific focus on the phylum *Bacillota*. We analyzed the diversity, persistence of ethanol-resistant bacteria, and sporulation frequency of ethanol-resistant *Bacillota* within and between nine healthy male adults sampled fortnightly over a six-month period.

## Methods

### Study design

The study included nine healthy, non-related and non-cohabitating male participants, aged 25 – 43, with no gastrointestinal infections or surgeries within three months prior to the study. Antibiotic use before or during the study was not an exclusion criterion. At the onset, each participant completed a questionnaire on health status and history, diet, and living conditions (Additional file 1). Stool samples were collected in approximately 14-day intervals and additionally at pre-determined events (dental visits, antibiotic use, travel, etc.) over a six-month period, with questionnaires filled out at each sample submission (Additional file 1). The participants used sterile collection containers for sampling and delivered the samples to the laboratory within a few hours of collection. If immediate delivery was not possible, samples were stored at -20 °C until transport. Upon receipt, each sample was homogenized, divided into five aliquots (one 10 µl inoculation loop per aliquot), and stored at -80 °C. Ethical approval was obtained from the National Medical Ethics Committee of the Republic of Slovenia (0120-280/2022/6).

### DNA isolation

DNA was extracted from two aliquots. The first aliquot was used to isolate total DNA from untreated fecal samples (bulk microbiota). The second aliquot was used to isolate DNA from ethanol-resistant bacteria, representing spores and other ethanol-resistant cell forms (ethanol treated sample). Bulk microbiota DNA was isolated using the PowerFecal Kit (Qiagen, Germany) following the manufacturer’s instructions and stored at -80 °C. A modified protocol from [12] was used for DNA isolation from the ethanol-resistant community. Stool aliquots were subjected to constant shaking in 70% ethanol for 24 hours, followed by washing with PCR-grade water. To remove free DNA and DNA in damaged cells, ethidium monoazide bromide (EMA) was added to a final concentration of 2.5 ng/µl. The samples were incubated for 5 min in the dark, followed by 10 min of exposure to an 8000-lumen light with ice cooling. Samples were washed again with PCR-grade water. DNA in the remaining ethanol treated community was isolated using the MasterPure Complete DNA and RNA Purification Protocol (LGC Biosearch Technologies, UK), with an additional bead-beating step (2 × 30 s at 7000 rpm, MagnaLyser (Roche, Switzerland), with 0.5 mm zirconia beads). The DNA obtained was treated overnight with RNase (5 mg/ml). The isolation efficiency of spores using the PowerFecal protocol was validated on stool samples with added spores of *Clostridioides difficile* (Additional file 2).

The DNA concentration of all samples was measured using the Quant-iT™ PicoGreen™ dsDNA Assay and RNA concentration using the Qubit™ RNA High Sensitivity Assay Kit (both Thermo Fisher Scientific, USA). The prepared DNA from the ethanol treated samples was additionally checked for protein contamination by measuring absorbance at 260 and 280 nm.

### Digital droplet PCR (ddPCR) for the 16S rRNA gene copy quantification

ddPCR was performed using EvaGreen technology (Bio-Rad QX200, USA) to quantify total bacterial load. ddPCR was conducted on all samples, using universal primers targeting the V3-V4 regions of the 16S rRNA gene 341F (5’CCTACGGGNGGCWGCAG-3’) and 806R (5’-GACTACHVGGGTATCTAATCC-3’) [19]. The reaction volume was 25 µl, consisting of 2.5 µl of diluted DNA, 12.5 µl of 2x EvaGreen Supermix (Bio-Rad), and 5 µl of each primer (1 µM). Droplets were generated using a QX200 Droplet Generator, following the manufacturer’s instructions. The thermal cycling conditions were as follows: enzyme activation at 95 °C for 3 min, followed by 40 cycles of denaturation at 95 °C for 30 s, annealing at 55 °C for 30 s, and elongation at 60 °C for 30 s, with a final elongation at 60 °C for 5 min, and signal stabilization at 4 °C for 5 min and 90 °C for 5 min. Quantification was performed using the QuantaSoft software (v7.4.1, Bio-Rad, USA), with a minimum threshold of 10,000 droplets per sample. The copy number of the 16S rRNA gene in each sample was calculated using the following Equation 1:

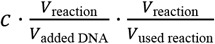 ·dilution factor·; 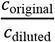 where *c* is concentration and *V* is volume.

Normalized abundance was calculated by multiplying the relative abundance of each OTU in the sample by the copy number of the 16S rRNA gene in that sample.

### Targeted sequencing of the 16S rRNA gene and sequence analysis

Bacterial community structure was determined by sequencing the V3-V4 region of the 16S rRNA gene using the same primers as for ddPCR (341F and 806R). Library preparation followed the 16S rRNA gene Metagenomic Sequencing Library Preparation protocol (Illumina, USA) with Q5® High-Fidelity DNA Polymerase (New England Biolabs, USA). Sequencing was performed using an Illumina NextSeq 2000 platform (2 × 300 cycles). Sequence reads were processed in mothur (v.1.48.0) according to the standard operating procedure for Illumina paired-end reads [20]. Reads were processed using the following criteria: reads were not allowed any ambiguous bases, and the maximum homopolymer length was set to 8 base pairs (bp); reads were aligned against the Silva reference alignment (Release 128) [21]; chimeras were identified using the UCHIME algorithm [22,23]; the classification of reads was performed using the RDP training set (v.9) [24], with a 0.80 bootstrap threshold value; sequences were clustered into operational taxonomic units (OTUs) at a 97% similarity cut-off. After quality filtering, we obtained an average depth of 270662.7 reads (min 61388, max 884743, not including negative control samples). OTUs represented by less than 0.0001% (62 reads) of the total number of reads or present in negative control samples were removed. Each sample was rarefied to 150,000 reads with rrarefy (R ‘vegan’ package) [25]. Samples with less than 150,000 reads were removed from further analysis (n=6, including both negative controls). Further statistical analysis and graphical representation were done in R (version 4.4.1) [26], and the code used to carry out the analysis presented in manuscript, along with notes showing command runs are available at: https://github.com/ursamiklavcic/longitudinal_amplicons. Sequence data is deposited in the SRA database, under the project name PRJNA1124620.

### Defining the ethanol-resistant or ethanol non-resistant community within bulk microbiota

Individual OTU within the bulk microbiota samples were determined as members of the ethanol-resistant community if the following criteria were met: 1) the OTU was present in both the bulk microbiota sample and the ethanol treated sample of the same stool sample, and 2) was found at a higher relative abundance in the ethanol treated sample in at least 5% of samples where criteria 1) were met. OTUs that were absent in all ethanol treated samples were considered ethanol non-resistant. OTUs with taxonomic assignment to the phylum *Bacillota* were placed in separate groups within the ethanol-resistant and ethanol non-resistant community. This was performed for two reasons. First, beta diversity comparisons could be skewed because of higher relative abundances of OTUs belonging to the phylum *Bacillota* as opposed to other taxa. Second, for the purpose of discussion, since all bacteria recognized as spore-forming in the human gut belong to this phylum, whereas the survival strategies of other phyla remain poorly understood [4–6,27,28]. We excluded OTUs that were detected in the ethanol-resistant community in less than 5% of the samples (determined uncertain; n = 83), and OTUs detected only in the ethanol treated samples (n = 141).

Beta diversity measures were calculated with ‘avgdist’ from the ‘vegan’ R package for Bray-Curtis dissimilarity and Jaccard distance [25]. The uneven sizes of the compared groups were addressed using bootstrapping (subsample size = 24, number of permutations = 999). Differences between group means were determined using the Wilcox rank-sum test.

### Sporulation frequency calculation and analysis of host, time, and OTU-specific influences

Sporulation frequency analysis was conducted on a subset of OTUs meeting two criteria: 1) classification within ethanol-resistant *Bacillota*, and 2) detection in ≥80% of bulk microbiota and ≥80% ethanol-treated samples (to ensure robust temporal coverage). For each eligible OUT (*i*) sporulation frequency was defined as the ratio between the normalized abundance of OTU in the ethanol treated sample (*e*_*i*_) and the normalized abundance of OTU in the bulk microbiota (*m*_*i*_), i.e., sporulation frequency = *e*_*i*_*/mi*. Here, the bulk microbiota includes sequences from both vegetative cells and ethanol-resistant forms, while the ethanol treated sample selectively enriches for spores and other ethanol-resistant cell forms (Additional file 2). Sporulation frequency reflects the fraction of an OTU population in a sporulated state at a given time point of an individual host. The reliability of this measure is supported by a strong correlation between relative and/or normalized abundance of OTU in both sample types, despite different isolation protocols (Additional file 2).

To assess whether the temporal order of sampling drives sporulation dynamics, we compared two scenarios for each host. First, where within-individual sporulation frequency was calculated for each OTU from chronologically ordered timepoints, and second, where time points were randomly shuffled prior to calculating the within-individual sporulation frequency for each OTU. Pearson correlation coefficients were then computed between ordered and reshuffled sporulation frequency values for each OTU-host pair to quantify the influence of temporal structure.

To characterize sporulation variability, we calculated the within-individual variance in sporulation frequency for each OTU and population-wide variance for this same OTU. Differences in the distribution of these variance values across hosts and OTUs were assessed using the Kruskal-Wallis test. To evaluate the degree of synchronization in sporulation dynamics across OTUs within individual hosts, we also computed pairwise Pearson correlation coefficients between sporulation frequencies of all OTUs within each individual.

## Results and discussion

Out of 2,431 OTUs detected in bulk microbiota samples, we determined 414 OTUs (17.0 %) as part of the ethanol-resistant community and 1,934 (79.5%) as part of the ethanol non-resistant community. Eighty-three OTUs (3.4%) failed to meet the criteria described in the Methods and were designated as undetermined regarding ethanol resistance. An additional 141 OTUs were detected only in ethanol treated samples. These were all detected at low relative abundance and were considered highly probable sampling artifacts and therefore excluded from further analysis, following an example from a previous study [11]. Compositional characteristics and temporal dynamics were subsequently analyzed on the above defined ethanol-resistant and non-resistant communities obtained from a 6-month longitudinal sampling of nine healthy males.

### Characteristics of ethanol-resistant and non-resistant communities

The majority of ethanol-resistant representatives were found in the phylum *Bacillota* (*n* = 320), accounting for 77.5% of the total richness in the ethanol-resistant community (Figure 1A). Within the phylum *Bacillota*, 17.7% of all detected OTUs exhibited ethanol resistance, predominantly from the genera *Clostridium* (n = 76), unclassified *Lachnospiraceae* (n = 49), unclassified *Ruminococcaceae* (n = 42), unclassified *Peptostreptococcaceae* (n = 13), and *Balutia* (n = 12). The remaining ethanol-resistant community included representatives from *Actinomycetota* (n = 42, 11%), *Pseudomonadota* (n = 21, 5.5%), and unclassified *Bacteria* (n = 14, 3.7%). In these groups, the proportion of ethanol-resistant OTUs was higher than expected, with 36.2%, 40.4%, and 19.7%, respectively, exhibiting this trait. Additionally, phyla with ten or fewer representatives collectively contributed 2.3% of the ethanol-resistant richness, most notably the biologically relevant genus *Akkermansia*. Ethanol treatment had the strongest elimination effect on the phylum *Bacteroidota*, with only 10 representatives remaining in the ethanol-resistant community, accounting for 0.5% of the total relative abundance.

**Figure 1.**
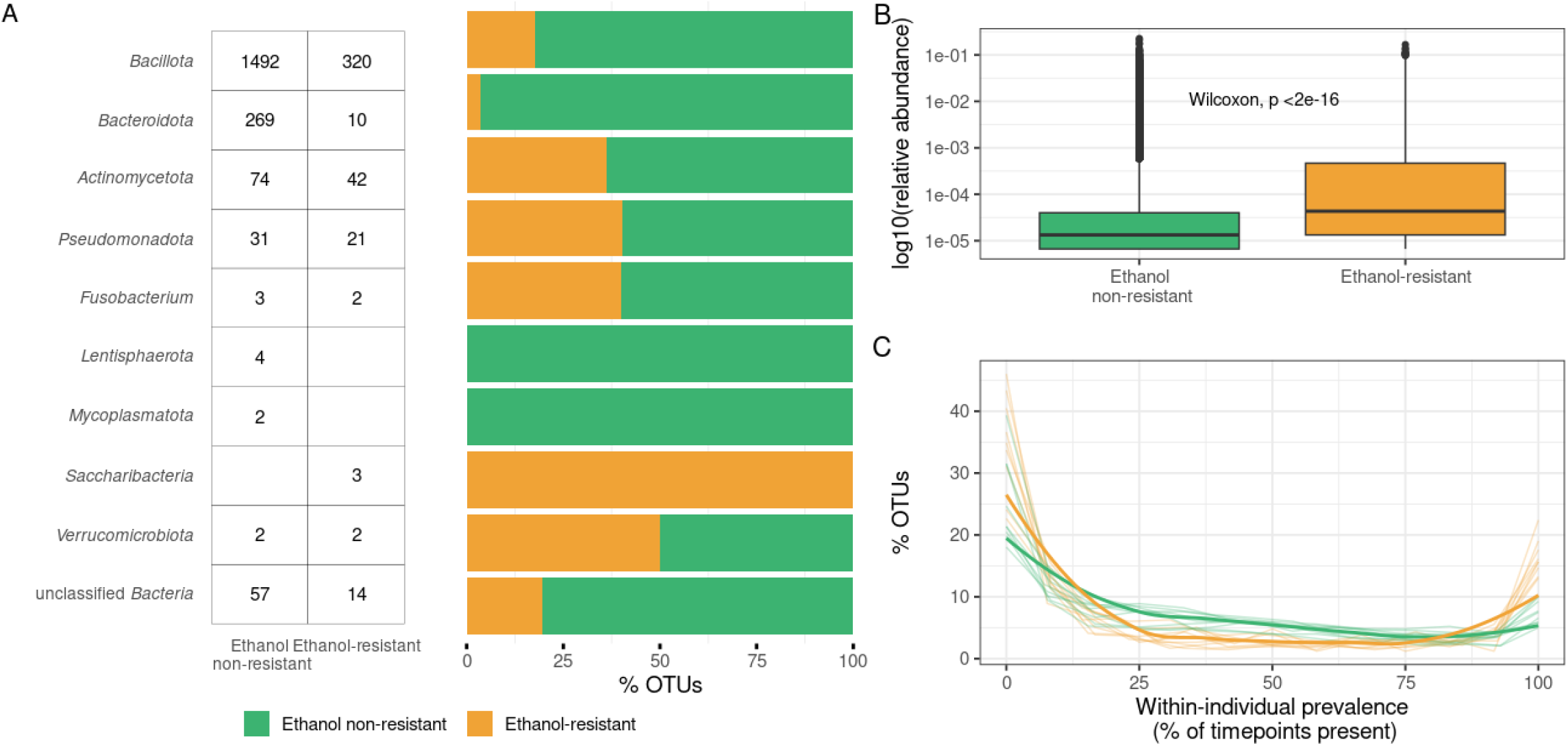
Characteristics of ethanol-resistant and ethanol non-resistant bacterial communities. A) Distributions of OTUs across phyla according to ethanol resistance. The table (left) shows the number of OTUs identified in each phylum for the ethanol non-resistant and ethanol-resistant communities, while the stacked bar plots (right) depict their relative proportions within the respective phyla, color-coded based on their ethanol non-resistant (green) or ethanol-resistant (yellow) classification. B) Comparison of relative abundances in the gut microbiota between OTUs determined as part of the ethanol-resistant or ethanol non-resistant community. C) Within-individual prevalence of OTUs across the six-month observational period, shown as the percentage of OTUs detected at different numbers of time points. Thin lines represent individual participants, and bold lines indicate the cohort median.

The high abundance of ethanol-resistant representatives within *Bacillota* was expected due to the widespread ability to sporulate within this phylum [5]. Most *Bacillota* representatives identified as ethanol-resistant in this study belong to genera in which sporulation has previously been confirmed through the detection of sporulation-associated gene clusters or *in vitro* sporulation induction [6,16,29]. In contrast, ethanol-resistant representatives, as well as the mechanisms underlying their resistance, are poorly described in other phyla. Previous studies have reported the detection of *Actinomycetota* representatives in ethanol or lysozyme treated samples [4,11,12]. While exospore formation was described in the families *Actinomycetaceae* and *Streptomycetaceae*, the mechanisms underlying ethanol resistance in other *Actinomycetota* families remain unclear. In our data, *Actinomycetaceae* accounted for only 0.4% of the relative abundance within the ethanol-resistant community, while *Streptomycetaceae* were not detected.

Members of the phylum *Pseudomonadota*, also commonly detected in the ethanol-resistant community, have previously been reported to adopt various dormant life strategies, such as persistence and the viable but non-culturable state [28]. A limitation of this study is the reliance on 16S rRNA amplicon sequencing, which does not provide information beyond taxonomic classification. Implementation of additional methodological approaches will be necessary to further elucidate the molecular basis of ethanol resistance in phyla other than *Bacillota*.

In general, we observed two notable characteristics distinguishing the ethanol-resistant community from the non-resistant community. Both were primarily driven by *Bacillota* OTUs due to their high abundance, though similar trends were also observed across other phyla. First, OTUs determined as ethanol-resistant exhibited significantly higher relative abundance within the gut microbiota compared to non-resistant OTUs (Wilcoxon rank-rum sum test, p < 0.001; Figure 1B). Second, ethanol-resistant OTUs were significantly more likely to be detected at more time points within an individual, particularly in more than 80% of samples collected over a six-month period, relative to non-resistant OTUs (Wilcoxon rank-sum test, p < 0.001; Figure 1C). These findings support the use of ethanol resistance as a proxy for identifying bacterial taxa employing dormancy mechanisms that contribute to long-term persistence in the human gut.

### Between and within-individual dynamics of ethanol-resistant and ethanol non-resistant communities over time

For the purpose of statistical robustness and clarity in discussion, we further divided our dataset into phylum *Bacillota* and other phyla (Figure 2). The ethanol-resistant communities exhibited significantly lower variability both between and within individuals compared to the non-resistant communities. This pattern was observed when implementing both unweighted (Jaccard distance, p < 0.001 for all comparisons; Figure 2A) and weighted beta diversity measures (Bray-Curtis, p < 0.001 for all comparisons; Additional file 3). However, the lower within-individual variability observed in ethanol-resistant communities was significantly more pronounced in the phylum *Bacillota* compared to other phyla (Wilcoxon rank-sum test, p < 0.001; Figure 2A). Our observation aligns with prior findings showing that lysozyme resistant taxa tend to have higher population-level prevalence and lower within-individual variability than non-resistant communities [11]. Moreover, a comparative genomic analysis has shown that bacterial species with more sporulation genes are significantly more prevalent in human populations, at least partially explaining the lower between-individual variability observed in this study [16].

**Figure 2.**
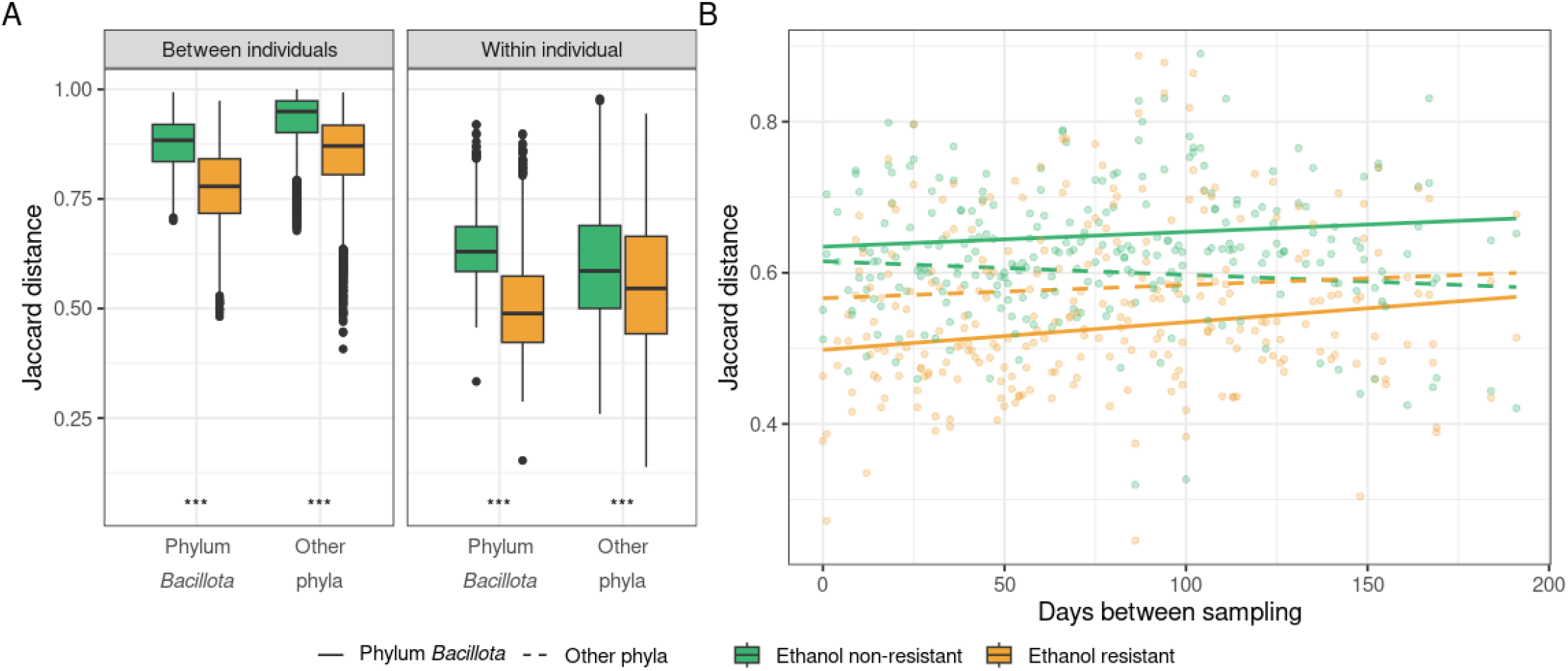
Dynamics of ethanol-resistant and ethanol non-resistant taxa between and within individuals. A) Boxplots showing between-individual (left) and within-individual (right) variability in community composition for ethanol-resistant (yellow) and non-resistant (green) communities. Variability was assessed using the Jaccard distance, calculated separately for OTUs from the phylum *Bacillota* and OTUs from other phyla. Jaccard distances were derived from randomly subsampled communities (n = 24, iterations = 999), with the median distance used for visualization and statistical comparisons. Distributions were compared using the Wilcoxon rank-sum test. B) Community variability as measured by Jaccard distance plotted against the number of days between sampling time points. Data are presented separately for phylum *Bacillota* (solid line) and other phyla (dashed line) and color-coded according to ethanol resistance. The correlation between time interval and Jaccard distance was assessed using Pearson’s correlation test. Statistical significance is indicated as *** for p < 0.001.

Spore persistence within the human gut was previously assessed using *in vitro* and murine gut models that have shown *C. difficile* spores bind to biofilms and consequently evade washout [30,31]. Their ability to persist is further supported by recurring infections with the same strain [32]. However, similar mechanisms of spore persistence have not been demonstrated for other species beyond *in vitro* biofilm formations [33]. In our study, we observed a modest but significant increase in temporal variability within ethanol-resistant *Bacillota* based on Jaccard distance (Pearson correlation, r = 0.19, p = 0.038, Figure 2B) and a borderline significant correlation when using Bray-Curtis dissimilarity (Pearson correlation, r = 0.169, p = 0.051, Additional file 3). In contrast, no significant temporal changes were observed in other communities (Figure 2B).

Our findings provide additional experimental evidence that spore formation in phylum *Bacillota* contributes to both increased transmission between individuals and long-term persistence within the host. In contrast, ethanol resistance mechanisms in non-*Bacillota* phyla appear to play a more prominent role in facilitating transmission, with a lesser impact on within-host persistence. A limitation of our study is the use of 16S rRNA amplicon sequencing, which does not provide taxonomic resolution beyond the genus. Consequently, we were unable to distinguish whether temporal changes in taxa detection reflected strain replacement or the coexistence of multiple strains [34]. This distinction is particularly relevant given previous reports of high strain-level retention in the gut over short time scales and limited strain sharing between individuals [35].

### Sporulation frequency of ethanol-resistant *Bacillota* in the human gut shows synchronized temporal variation and is influenced by the individual host

We calculated sporulation frequency for OTUs belonging to the ethanol-resistant community and phylum *Bacillota* (see Methods) that were present in at least 80% of all samples (n = 20). Sporulation frequency was defined as the ratio between normalized abundance of an OTU in ethanol-treated samples (spores) and its normalized abundance in bulk microbiota (vegetative cells + spores). Sporulation frequency was subsequently examined to determine whether it was primarily influenced by host-specific factors, temporal variation, or the identity of individual OTUs (Figure 3).

**Figure 3.**
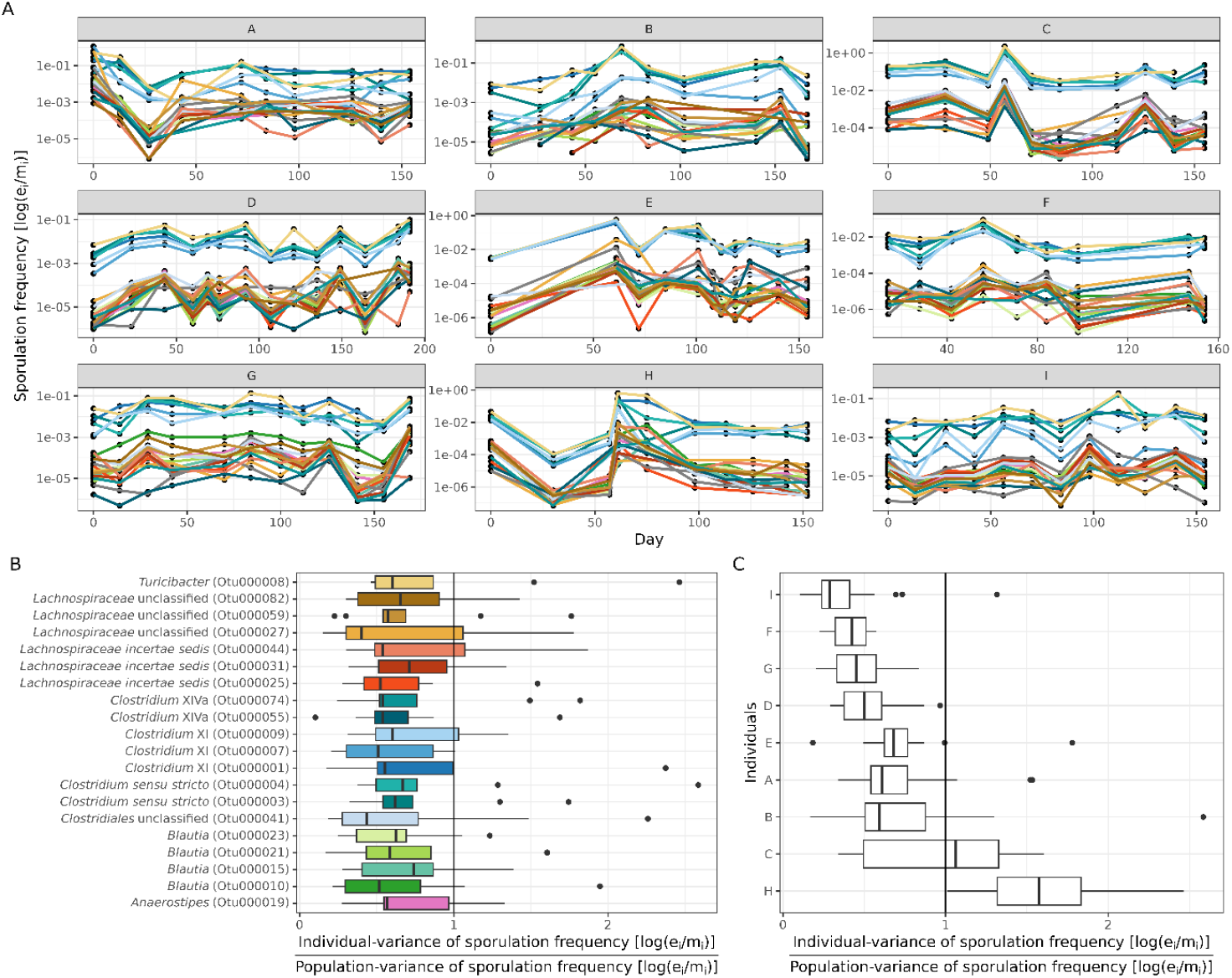
Influence of individual, time or OTU identity on sporulation frequency. The analysis includes only highly prevalent OTUs belonging to ethanol-resistant *Bacillota* (present in > 80% of samples, n = 20). A) Sporulation frequency of each OTU over the study duration, shown separately for each participant. Sporulation frequency was calculated as the log10-transformed ratio of normalized OTU abundance and ethanol-treated samples (e_*i*_) and bulk microbiota samples (m_*i*_). B) Ratio of within-individual to population-level variance in sporulation frequency, plotted by OTU. C) Ratio of within-individual to population-level variance in sporulation frequency, plotted by study participant. The vertical line (x = 1) indicates where within-individual (OTU or participant) variance equals the population-wide variance.

The OTUs in the above defined subset exhibited individual-specific and highly synchronized, temporal fluctuations in sporulation frequency across taxonomically distinct taxa (Figure 3A). A lack of correlation between the original and temporally shuffled data (Pearson correlation: 0.03, p = 0.74) suggests that sporulation frequency was influenced by the current state of the gut environment in each individual. These individual-specific patterns were also supported by significant positive correlations in sporulation frequency among OTUs within the same individual (Additional file 4). Furthermore, the within-individual variance in sporulation frequency for an OTU was significantly lower than the population-wide variance (Kruskal-Wallis, p = 0.96; Figure 3B). Together, these results suggest that host-specific gut environments have a greater impact on sporulation frequency than OTU identity.

Among nine study participants, two individuals (H and C) displayed higher variance in sporulation frequency compared to the population mean (Kruskal test, p < 0.001; Figure 3C). These deviations coincided with antibiotic use between day 71 and 84 (antibiotic amoxicillin; individual H) and 5-day trip abroad during the second month of the study (individual C). These changes also coincided with shifts in alpha diversity in bulk microbiota, decreasing in individual H and increasing in individual C. While other participants also experienced travel (A, B, D) or antibiotic use (antibiotic doxycycline; E) during the study, these events did not notably affect their sporulation frequency.

Our observations align with previous studies demonstrating that external stimuli, such as dietary or environmental changes, impact the gut microbiota in a host specific manner [36]. Additionally, our results suggest that the sporulation frequency of ethanol-resistant *Bacillota* varies between individuals and fluctuates over time within an individual but remains synchronized across different taxa within the same host (Figure 3). *In vitro* studies have shown that sporulation and germination in individual bacterial species can be triggered by environmental stress, metabolic depression, inter-bacterial signaling, or viral threats [29,37,38]. Identifying specific *in vivo* signals driving sporulation and germination was beyond the scope of this study. Nevertheless, our findings suggest that the spore-forming community in the human gut responds collectively to shared, yet unidentified, environmental, or physiological signals, rather than to intrinsic signals from individual species.

## Conclusions

This study provides new insights into the structure and dynamics of ethanol-resistant taxa within the human gut microbiota, with a particular emphasis on spore-forming *Bacillota*. We identified a distinct ethanol-resistant community from eight bacterial phyla. Compared to the non-resistant community, ethanol-resistant taxa showed greater similarity both within and between individuals, an effect especially pronounced among *Bacillota*. Moreover, the sporulation frequency of selected highly prevalent *Bacillota* displayed temporal and host-specific patterns, with synchronous fluctuations observed across different taxa within the same individual. Collectively, our findings suggest that sporulation is a dynamic, community-wide process shaped primarily by the host environment.

## Supporting information

Additional file 2, Justification of sporulation frequency calculation

Additional file 3, Community dynamics between and within individuals using Bray-Curtis dissimilarity

Additional file 4, Correlation of OTUs sporulation frequency within an individual

Additional file 1, Participant questionnaires

## List of abbreviations

PCR: polymerase chain reaction
OTU: operational taxonomic unit
ddPCR: digital droplet PCR

## Additional file information

Additional file 1, additional_file1.pdf, Participant questionnaires

Additional file 2, additional_file2.pdf, Justification of sporulation frequency calculation

Additional file 3, additional_file3.pdf, Community dynamics between and within individuals using Bray-Curtis dissimilarity

Additional file 4, additional_file4.pdf, Correlation of OTUs sporulation frequency within an individual

## Declarations

### Ethics declaration

This study was conducted in compliance with the recommendations of the Declaration of Helsinki and approved by the Commission of the Republic of Slovenia for Medical Ethics (no. 0120-280/2022/6).

### Consent for publication

All participants have consented to participate in the study and understood the data shared.

### Availability of data and materials

The datasets generated and analyzed during the study are available in the GitHub repository, at www.github.com/ursamiklavcic/longitudinal_amplicons. Raw sequences are deposited in SRA database, under the project name PRJNA1124620.

### Authors’ contributions

MR contributed to study concept and design, funding acquisition and revision of manuscript. SJ contributed to study concept and design. AM, WRS and JG contributed to data analysis, data interpretation and revision of manuscript. UM contributed to study design and concept, sample and data acquisition, data analysis and interpretation, drafting and revising the manuscript. All authors have read and approved the manuscript and accept personal accountability for their work.

