## Additional file 2, Justification of sporulation frequency calculation for "Longitudinal dynamics of ethanol-resistant microbes and sporulation in the human gut"

### DNA isolation efficiency from spores using the PowerFecal protocol and *Clostridioides difficile* spores

To confirm that we isolated both vegetative cells and spores with bulk microbiota protocol, we added *Clostridioides. difficile* spores to an otherwise *C. difficile* negative stool sample and measured the absolute amount of the *C.difficile*-specific 16S rRNA amplicon with ddPCR after isolation of DNA. *C. difficile* (strain VPI10463) was cultured from spore stock solution and plated confluently on three Columbia agar plates with 5% added horse blood (COH) plates (Biomérieux, France). After five days of anaerobic incubation at 37 °C, spores and cells were harvested and washed five times with PCR-grade water. A 1 ml aliquot of washed spores was treated with EMA (2.5 ng/µl), sample was incubated for 5 minutes in the dark, followed by 10-minute exposure to 8000-lumen light with ice cooling. Fresh stool samples were homogenized and aliquoted (~100 mg) into PowerFecal bead-beating tubes. DNA was isolated from fresh stool samples and from samples spiked with spores at various dilutions (non-diluted, $10^{-1}$, $10^{-3}$, and $10^{-4}$), in triplicates. The same spore dilutions were also plated on COH agar to determine spore concentration with CFU count. Digital droplet PCR (ddPCR) was performed using EvaGreen technology (Bio-Rad QX200) for the quantification of *C. difficile*. ddPCR was conducted using *C. difficile*-specific primers targeting the 16S rRNA gene F(5' CCATCCTGTACTGGCTCACCT-3') and R(5'- TTGAGCGATTTACTTCGGTAAAGA-3') [1]. The reaction volume was 25 µl, consisting of 2.5 µl of diluted DNA, 12.5 µl of 2x EvaGreen Supermix (Bio-Rad), 0.5 µl of each 10 µM primer, and 9 µl of PCR-grade water. Droplets were generated using the QX200 Droplet Generator, following the manufacturer's instructions. The thermal cycling conditions for *C. difficile* 16S rRNA were enzyme activation at 95 °C for 5 minutes, followed by 40 cycles of denaturation at 95 °C for 30 seconds, annealing and elongation at 60 °C for 60 seconds, and signal stabilization at 4 °C for 5 minutes and 90 °C for 5 min. Absolute quantification was performed using QuantaSoft software (v7.4.1, Bio-Rad), with a minimum threshold of 10,000 droplets per sample. We calculated the copy number of 16S rRNA gene in each sample with Equation 1:

$c \cdot \frac{V_{\text{reaction}}}{V_{\text{added DNA}}} \cdot \frac{V_{\text{reaction}}}{V_{\text{used reaction}}} \cdot \text{dilution factor} \cdot \frac{c_{\text{original}}}{c_{\text{diluted}}}$; where $c$ is concentration and $V$ is volume. Copy number of the *C. difficile*-specific 16S rRNA gene in the samples was compared to CFU counts of spores.

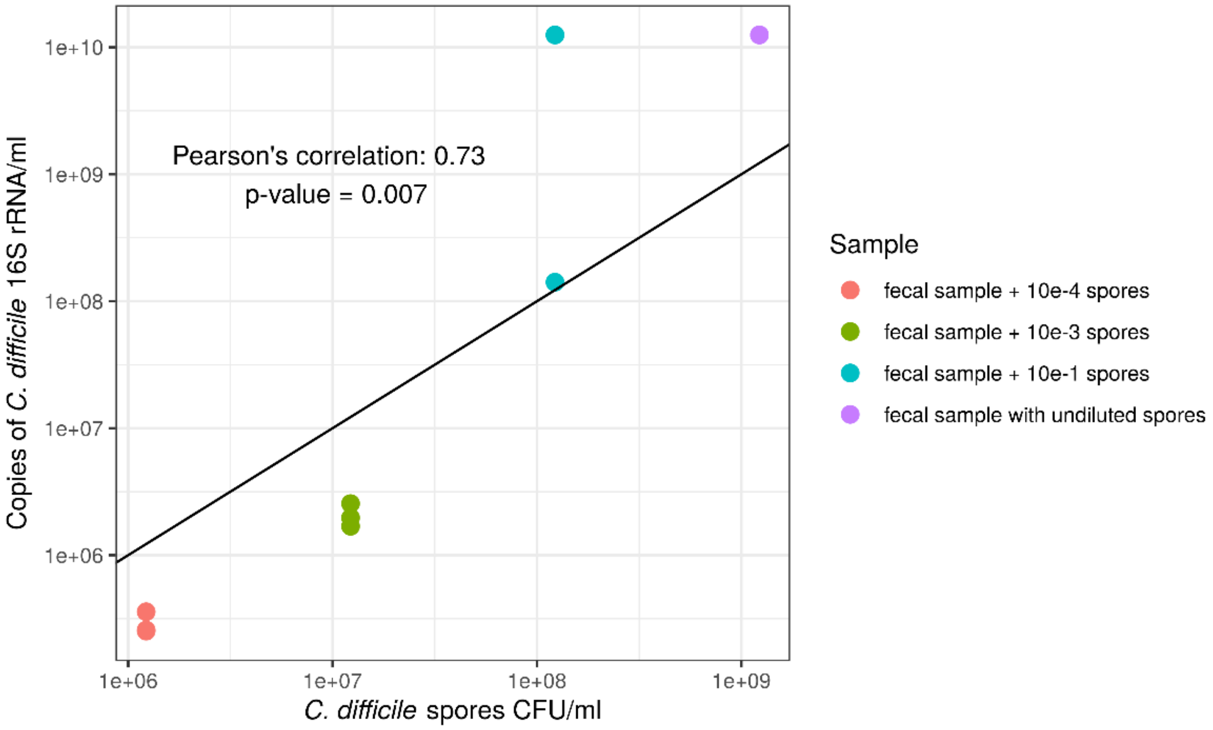


**Additional Figure 1. Efficiency of *C. difficile* spore isolation with PowerFecal protocol compared to *C. difficile* spore CFU count on COH agar plates.** The copy number of C. difficile-specific amplicon was correlated to CFU count of spores after cultivation (Pearsons correlation value = 0.73, p-value = 0.007). Ethanol shock is a known protocol for removal of vegetative cells as 70% ethanol damages the cell wall of all known bacteria in the human gut but does not damage spores and other ethanol resistant cell forms [2].

**Correlation of relative and/or normalized abundance in ethanol treated samples and bulk microbiota samples**

Comparison of the relative abundance in the bulk microbiota sample and ethanol treated sample fir ethanol resistant OTUs revealed that these two quantities are, despite different isolation protocols positively correlated (Additional Figure 2). The same holds true for normalized abundances (Additional Figure 3). We could thus compare the proportion of active cells that are also creating spores. The ratio gives us the information on the sporulation frequency of each OTU determined as endospore forming.

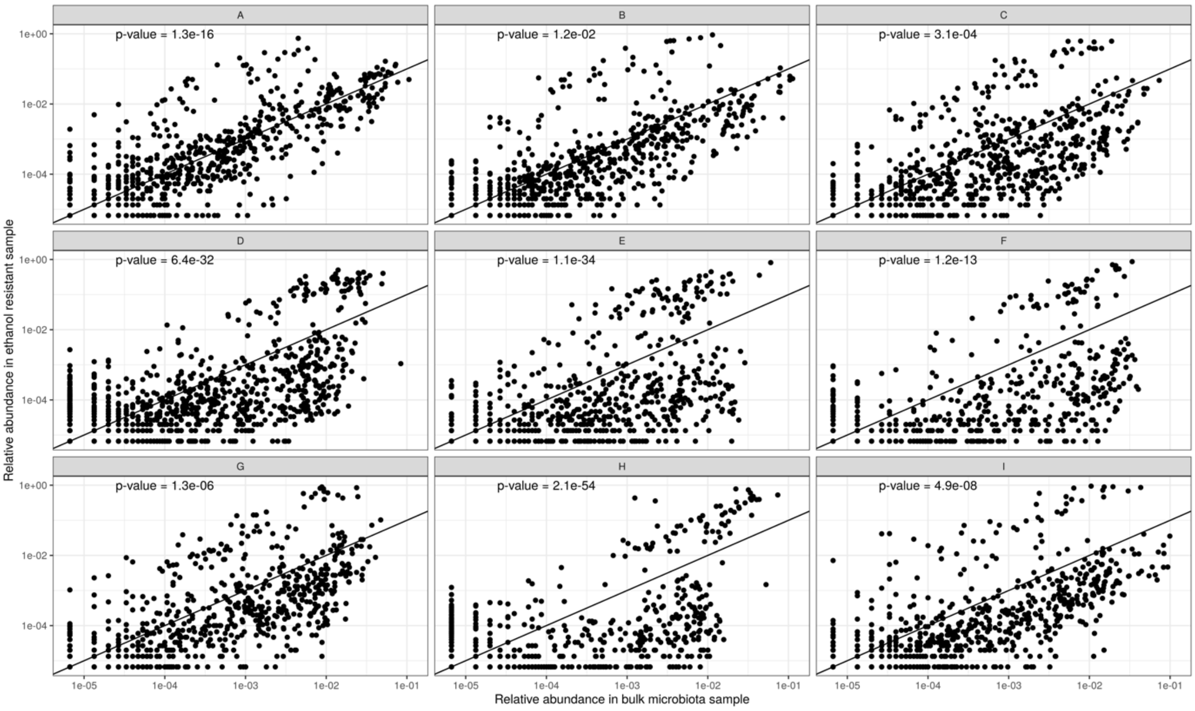


**Additional Figure 2.** Relative abundance in ethanol treated sample and relative abundance in microbiota sample are positively correlated (p-value for each individual Pearson's correlation is plotted in the top left corner of the plot).

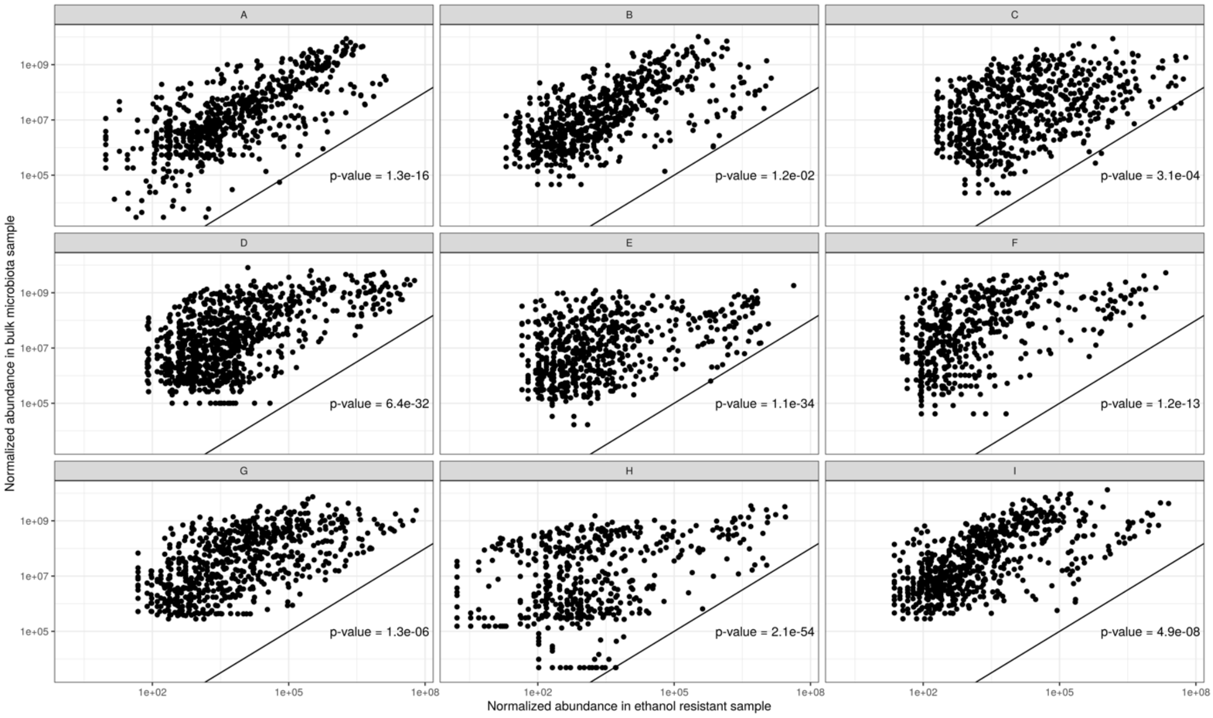


**Additional Figure 3.** Normalized abundance in ethanol treated sample and normalized abundance in microbiota sample are positively correlated (p-value for each individual Pearson's correlation is plotted in the bottom right corner of the plot).
