## Additional file 3, Community dynamics between and within individuals using Bray-Curtis dissimilarity for "Longitudinal dynamics of ethanol-resistant microbes and sporulation in the human gut"

**Additional file 3: Bray-Curtis dissimilarity comparison between, within individual and within individuals through time & results for Wilcox rank sum test and Pearon correlation**

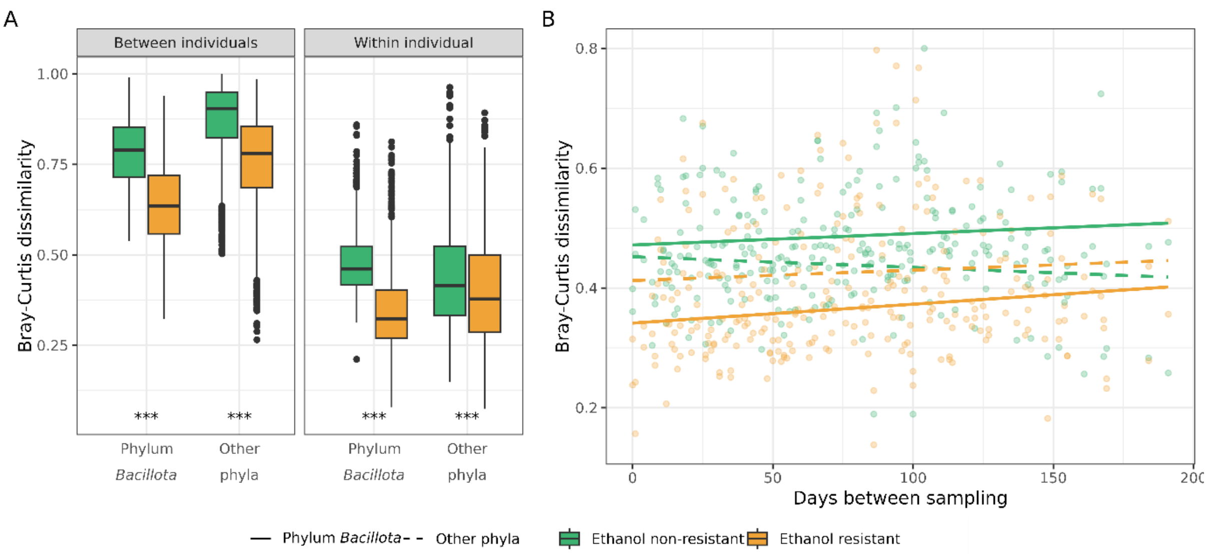


**Additional Figure 4. Dynamics of ethanol-resistant and ethanol non-resistant taxa between and within individuals.** A) Boxplots showing between-individual (left) and within-individual (right) variability in community composition for ethanol-resistant (yellow) and non-resistant (green) communities. Variability was assessed using the Bray-Curtis dissimilarity, calculated separately for OTUs from the phylum *Bacillota* and OTUs from other phyla. Bray-Curtis dissimilarity was derived from randomly subsampled communities (n = 24, iterations = 999), with the median distance used for visualization and statistical comparisons. Distributions were compared using the Wilcoxon rank-sum test. B) Community variability as measured by Bray-Curtis dissimilarity plotted against the number of days between sampling time points. Data are presented separately for phylum *Bacillota* (solid line) and other phyla (dashed line) and color-coded according to ethanol resistance. The correlation between time interval and Jaccard distance was assessed using Pearson's correlation test. Statistical significance is indicated as *** for $p < 0.001$.

**Additional Table 1.** Wilcox rank sum test between all fractions inter and intra individuals.

| | | p-value | | | |
|---|---|---|---|---|---|
| | | Bray-Curtis dissimilarity | | Jaccard distance | |
| Fraction 1 | Fraction 2 | Within individual | Between individuals | Within individual | Between individuals |
| Ethanol resistant Bacillota | Ethanol non-resistant Bacillota | < 0.0001 | 2.7e-194 | 8.2e-200 | < 0.0001 |
| Other ethanol non-resistant taxa | Ethanol non-resistant Bacillota | < 0.0001 | 1.5e-31 | 6.4e-38 | < 0.0001 |
| Other ethanol resistant taxa | Ethanol non-resistant Bacillota | 3.5e-18 | 2.2e-63 | 6.6e-65 | 7.1e-25 |
| Other ethanol non-resistant taxa | Ethanol resistant Bacillota | < 0.0001 | 4.2e-65 | 1.3e-60 | < 0.0001 |
| Other ethanol resistant taxa | Ethanol resistant Bacillota | < 0.0001 | 9.8e-18 | 2.6e-19 | < 0.0001 |
| Other ethanol resistant taxa | Other ethanol non-resistant taxa | < 0.0001 | 3.2e-11 | 6.9e-09 | < 0.0001 |

**Additional Table 2.** Pearson correlation values and p-values between median distance and time spans between an individual's fractions. Significance is denoted as: * 0.05 – 0.01, **0.01 – 0.001 and *** < 0.001.

| | Bray-Curtis dissimilarity | | Jaccard distance | |
|---|---|---|---|---|
| Fraction | Pearsons correlation | p-value | Pearsons correlation | p-value |
| Ethanol resistant Bacillota | 0.161 | 0.0624 | 0.180 | 0.0369 * |
| Ethanol non-resistant Bacillota | 0.135 | 0.120 | 0.158 | 0.0676 |
| Other ethanol resistant taxa | 0.068 | 0.429 | 0.0683 | 0.433 |
| Other ethanol non-resistant taxa | - 0.078 | 0.366 | -0.105 | 0.226 |
