## Additional file 4, Correlation of OTUs sporulation frequency within an individual for "Longitudinal dynamics of ethanol-resistant microbes and sporulation in the human gut"

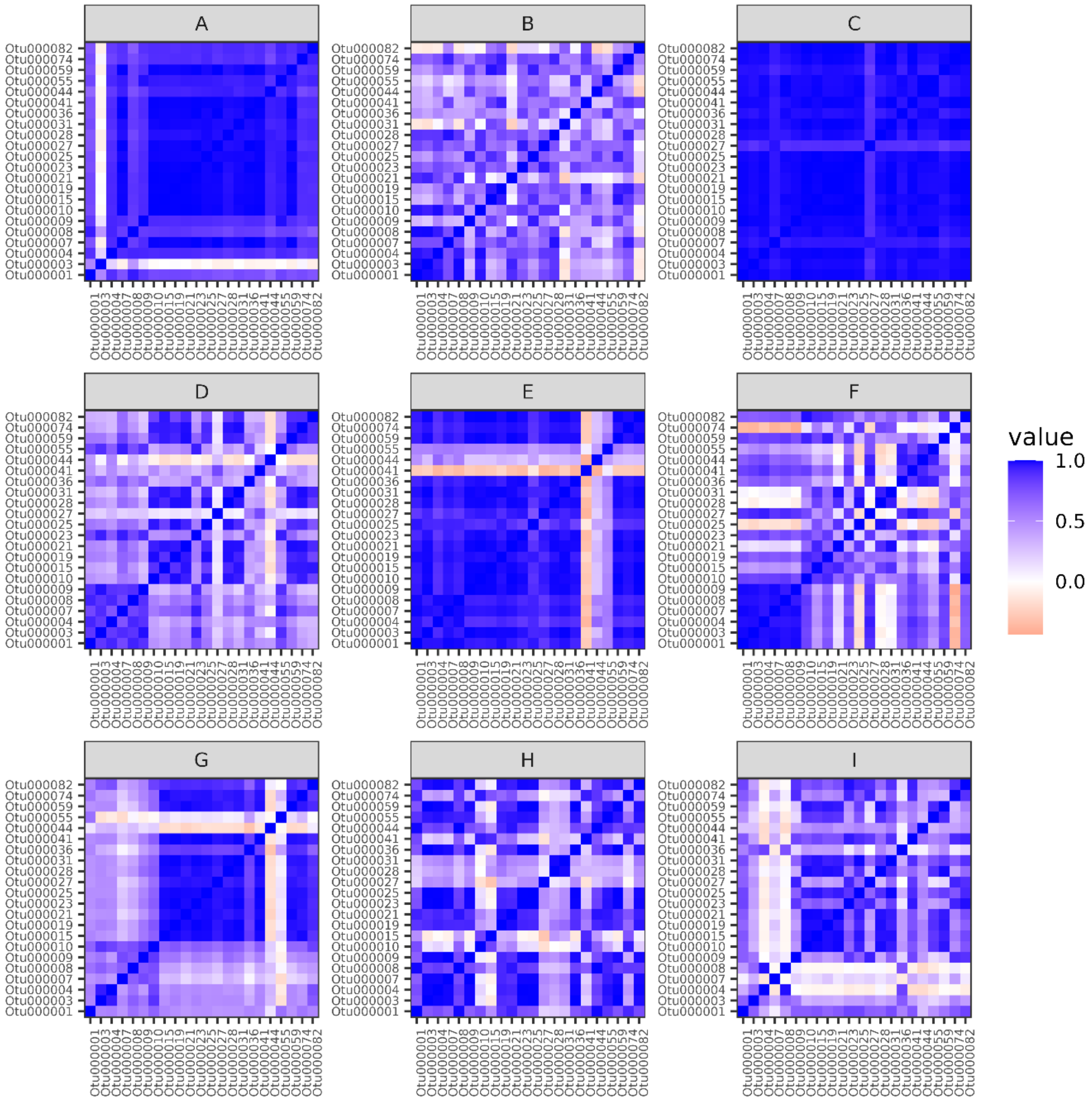


**Additional Figure 5.** Correlation of sporulation frequency between OTUs within individuals. Mostly OTUs of an individual are correlated within an individual, which point to host specific sporulation frequency.
