## Additional file 1, Participant questionnaires for "Longitudinal dynamics of ethanol-resistant microbes and sporulation in the human gut"

**Additional file 1. Questionnaires for participants.** Original questionnaires were in Slovene.

### Questionnaire 1: Before entering the study

As part of the Longitudinal Stability of the Gut Microbiota study for the PhD of Ursa Miklavcic, please fill in the questions below, as these questions, together with your samples, will give a comprehensive insight into the stability of your gut microbiota over a six-month period.

You are completing the questionnaire under your unique code so that we can protect your anonymity and confidentiality.

Thank you very much for your participation.

If you have any further questions, please contact me by e-mail: _________or by phone ________.

Ursa Miklavcic

**Code:**

---

**PLEASE FILL IN!**

Sex: ☐ M ☐ F    Age (years): _________________

Height: _________    Weight: ___________

Postal code: _________________

We will remind you a few days before the sample is submitted, so you have given your consent to the use of your personal data. Please provide us with your telephone number: ______________________ or email: _________________________.

---

Please answer the following questions below.

Eating habits:

- ☐ No specifics
- ☐ Vegi
- ☐ Vegan
- ☐ Gluten-free
- ☐ Lactose-free
- ☐ No-sugar
- ☐ Paleo diet
- ☐ Keto diet
- ☐ Only fresh food
- ☐ Other: _______________________

Do you have a medically confirmed allergy to:

☐ Gluten

- ☐ Lactose
- ☐ Peanuts
- ☐ Other nuts
- ☐ Eggs
- ☐ Cochineal
- ☐ Other: ____________________

Do you take any supplements?

- ☐ No
- ☐ Yes:
  - ☐ Omega 3
  - ☐ Vitamin D
  - ☐ Vitamin B12
  - ☐ Complex vitamins B
  - ☐ Vitamin C
  - ☐ Q10 complex
  - ☐ Other: ____________

Physical activity

- ☐ Occasionally (less than 1x per week)
- ☐ Recreational 1x per week
- ☐ Recreational more than 1x per week
- ☐ Active athlete

Regular smoker (several times a week)? ☐ No ☐ Yes

Did you take antibiotics in the last 3 months?

- ☐ No
- ☐ Yes
- ☐ Don't know

Have you ever had appendectomy in the past? ☐ No ☐ Yes

Have you ever had gallstone surgery in the past? ☐ No ☐ Yes

Have you had any other gastrointestinal surgery in the past? ☐ No ☐ Yes

Have you been hospitalized for more than two days in the last three months? ☐ No ☐ Yes

Have you had any gastrointestinal infections (vomiting and/or diarrhea) in the past 3 months?
☐ No ☐ Yes

Please mark the presence of any of the underlying gastrointestinal diseases (multiple answers are possible):

☐ Irritable bowel syndrome (IBS)

☐ Chron's disease

☐ Ulcerative colitis

☐ Common/persistent non-specific gastrointestinal problems (e.g. flatulence, pain, cramps, wind, nausea, etc.)

☐ Other: ______________________________

If you have ticked common/persistent non-specific problems which?

☐ Bloating

☐ Pain in the abdomen

☐ Cramps

☐ Gases

☐ Sickness

☐ Other: ______________________________

How would you assess your digestion?

☐ Regular (daily most of the time)

☐ Occasional problems with constipation (constipated maximum 2 days per week)

☐ Major constipation problems (constipated more than 2 days a week)

How many people do you live with?

☐ 1

☐ 2

☐ 3 and more

I live in a:

- ☐ house
- ☐ apartment
- ☐ do not want to disclose.

I live in ☐ rural or ☐ urban environment.

Thing you wish for us to know:

### Questionnaire 2: Submitted with each stool sample submission

As part of the Longitudinal Stability of the Gut Microbiota study for the PhD of Ursa Miklavcic, please fill in the questions below, as these questions, together with your samples, will give a comprehensive insight into the stability of your gut microbiota over a six-month period.

You are completing the questionnaire under your unique code so that we can protect your anonymity and confidentiality.

Thank you very much for your participation. If you have any further questions, please contact me by e-mail.

Ursa Miklavcic

**Date:** **Code:**

---

**<u>The questions below refer to the past 14 days.</u>**

Eating habits:

- ☐ No specifics
- ☐ Vegi
- ☐ Vegan
- ☐ Gluten-free
- ☐ Lactose-free
- ☐ 1 to more days without food
- ☐ No-sugar
- ☐ Other: _______________

Did you take antibiotics?

- ☐ No
- ☐ Yes, which:

_________________________, how many days ________________?

Did you take probiotics?

- ☐ No
- ☐ Yogurt with added probiotics
- ☐ Other: ________________________

Did you take prebiotics?

- ☐ No
- ☐ Yes, which _____________________

Did you take any medication under supervision of your doctor?

- ☐ No
- ☐ Yes, which _____________________

Did you take any supplements?

- ☐ No
- ☐ Yes, which _____________________

- ☐ More sugar

How would you describe the levels of your stress ☺ 1 2 3 4 5 ☹

Did any of the following events happen?
- ☐ Moving house
- ☐ Extreme physical activity (irregular from your baseline)
- ☐ Travel abroad, where: ____________
- ☐ Travel in Slovenia (more than 3 days)

Describe your level of activity in the past 14 days:
at least 30 min of moderate activity (walk, yoga, walking to work, playing with children, ..)
- ☐ 1-2x
- ☐ 2-3x
- ☐ 4x or more

Active (biking, fitness, hiking, …)
- ☐ 1-2x
- ☐ 2-3x
- ☐ 4x ali več

Did you visit the dentist?
- ☐ No
- ☐ Yes

Have you had an invasive body procedure (surgery or examination where your body has been interfered with, tattoo, piercing, etc.)?
- ☐ No
- ☐ Yes, what _________________

Have you been vaccinated in the past 14 days?
- ☐ No
- ☐ Yes, what vaccine ___________________?

Please, asses your stool, type ______________.

| | | |
|---|---|---|
| *Type 1* | 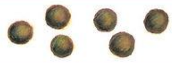 | Separate hard lumps, like nuts (hard to pass) |
| *Type 2* | 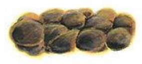 | Sausage-shaped but lumpy |
| *Type 3* | 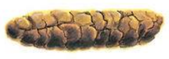 | Like a sausage but with cracks on its surface |
| *Type 4* | 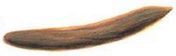 | Like a sausage or snake, smooth and soft |
| *Type 5* | 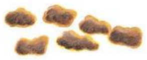 | Soft blobs with clear-cut edges (passed easily) |
| *Type 6* | 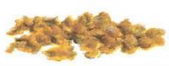 | Fluffy pieces with ragged edges, a mushy stool |
| *Type 7* | 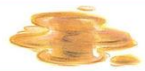 | Watery, no solid pieces ENTIRELY LIQUID |
